# Top-Down Beta Oscillatory Signaling Conveys Behavioral Context to Primary Visual Cortex

**DOI:** 10.1101/074609

**Authors:** Craig G. Richter, Richard Coppola, Steven L. Bressler

## Abstract

Top-down modulation of sensory processing is a critical neural mechanism subserving a number of important cognitive roles. Principally, top-down influences appear to inform lower-order sensory systems of the current ‘task at hand’, and thus may convey behavioral context to these systems. Accumulating evidence indicates that top-down cortical influences are carried by directed interareal synchronization of oscillatory neuronal populations. An important question currently under investigation by a number of laboratories is whether the information conveyed by directed interareal synchronization depends on the frequency band in which it is conveyed. Recent results point to the beta frequency band as being particularly important for conveying task-related information. However, little is known about the **nature** of the information conveyed by top-down directed influences. To investigate the information content of top-down directed beta-frequency influences, we measured spectral Granger Causality using local field potentials recorded from microelectrodes chronically implanted in visual cortical areas V1, V4, and TEO, and then applied multivariate pattern analysis to the spatial patterns of top-down spectral Granger Causality in the visual cortex. We decoded behavioral context by discriminating patterns of top-down (V4/TEO → V1) beta-peak spectral Granger Causality for two different task rules governing the correct responses to visual stimuli. The results indicate that top-down directed influences in visual cortex are carried by beta oscillations, and differentiate current task demands even before visual stimulus processing. They suggest that top-down beta-frequency oscillatory processes may coordinate the processing of sensory information by conveying global knowledge states to early levels of the sensory cortical hierarchy independently of bottom-up stimulus-driven processing.

## Introduction

Perception is not driven solely by sensory stimulation. Rather, endogenous processing actively modulates and routes sensory input based on prior knowledge, adapts perception to satisfy task demands, and solves ambiguities in the sensory stream (Engel et al. 2001; Gilbert and Sigman 2007; Wang 2010). Mounting evidence indicates that oscillatory activity conveys information between visual cortical areas (Fries 2005; Bressler and Richter 2015; Fries 2015). Anatomical studies show that cortical areas are linked via unique patterns of cortical projections and terminations between the cortical laminae that define the cortical hierarchy (Felleman and van Essen, 1991; Hilgetag et al. 1996; Markov et al. 2014). Recent studies have revealed that information transfer across the cortical hierarchy occurs in unique frequency regimes, with gamma frequency rhythms subserving bottom-up (feedforward) information transfer, while beta rhythms mediate transfer in the reverse (top-down) direction (Bressler et al. 2007, Buschman and Miller 2007; Bosman et al. 2012; van Kerkoerle et al. 2014; Bastos et al. 2015; Richter et al. 2016; Michalareas et al. 2016). Furthermore, top-down beta frequency influences may directly affect stimulus-related processing, as indicated by recent studies demonstrating that the magnitude of top-down beta-frequency rhythms is increased to the hemisphere representing an attended stimulus, resulting in enhanced bottom-up processing of the attended stimulus (Bosman et al. 2012; Bastos et al. 2015; Richter et al. 2016). Top-down beta-frequency synchronization may play a general role in behavior by conveying moment-to-moment task demands of the organism to lower level sensory systems in order to maintain global knowledge states (Engel and Fries 2010; Bressler and Richter 2015). Specifically, top-down beta rhythms may mediate the interaction between endogenously generated top-down information, variously described as hypotheses, prior knowledge, or attentional locus, and stimulus-generated bottom-up information (Bastos et al. 2012; Bressler and Richter 2015, Richter et al., 2016). Consequently, we hypothesize that behavioral context is encoded in the pattern of top-down beta synchronization in visual cortex.

We tested this hypothesis in two macaque monkeys performing a visual discrimination task (Figure 1), in which the behavioral context was determined by the task rule governing the correct response to each visual stimulus, the rule being randomly varied across trial blocks. Local field potentials were chronically recorded from microelectrodes in primary visual cortex (area V1) and extrastriate areas V4 and TEO. Both top-down (V4/TEO-to-V1) and bottom-up (V1-to-V4/TEO) directed synchrony were quantified using spectral Granger Causality in a stationary time period when the animal could anticipate the visual stimulus, but before it was presented. In this way, top-down influences were isolated from any confounding effects of stimulus processing. We successfully decoded behavioral context at a level significantly exceeding chance (50%) in both monkeys (76% and 82%) by two-class multivariate pattern analysis, using the spatial pattern of prestimulus top-down beta synchrony directed from V4/TEO to V1 as the classification feature.

**Figure 1.**
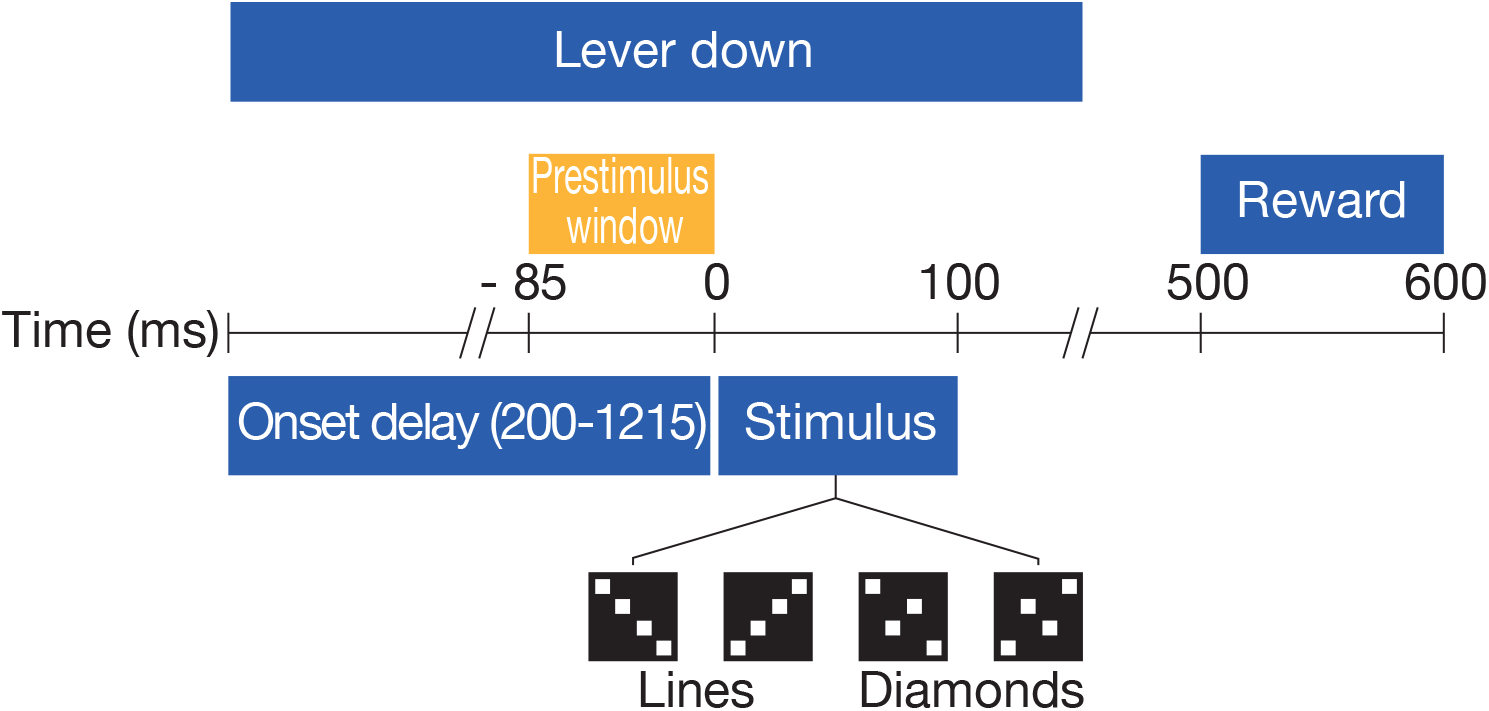
Task Structure. Task components of a Go trial are shown in blue, with the analysis window in gold. No-go trials followed the same event time course, except that for no-go trials the lever press was maintained throughout the trial, and there was no reward.

The results of this study indicate that the endogenous pattern of top-down beta influence from V4/TEO carries task-specific information to primary visual cortex (V1). We conclude from this that beta oscillations adaptively coordinate the processing of sensory information by conveying behavioral context to early levels of the sensory cortical hierarchy. We infer that behavioral context is conveyed from higher to lower levels of visual cortex by beta-frequency synchronized oscillations because pattern classification analysis successfully discriminated different visuomotor contingencies using higher-to-lower-level beta-frequency visual cortical influences as classification features. Our results thus support the notion that top-down beta-frequency oscillations play a general role in mediating the interaction between high-level cognitive processing and stimulus-related activity.

## Results

### Prestimulus Beta-Frequency Oscillatory Synchrony in Visual Cortex

Beta-frequency oscillations in V1 and V4/TEO were detected as spectral peaks near 16 Hz in the de-noised LFP power spectra (Figure 2a,b) computed during a brief prestimulus window. Generally, LFP time series are well described as stochastic processes, and spectral power peaks indicate the frequency and magnitude of narrow-band oscillatory activity in those processes. During the prestimulus period, the monkey had already pressed a lever (and maintained it in the depressed position) to begin the trial, and was awaiting well known visual stimuli. Prestimulus synchronization of the V1 and V4/TEO beta-frequency oscillations was observed as a peak in the average V1-extrastriate coherence spectrum at 17 Hz (Figure 2c). Coherence is a linear approximation of the total interdependence *F*_*xy*_ for values in the range (<~0.2) normally encountered physiologically (see Supplementary Figure 2). A peak in the coherence spectrum indicates narrow-band synchronization between the two processes used to derive it.

**Figure 2.**
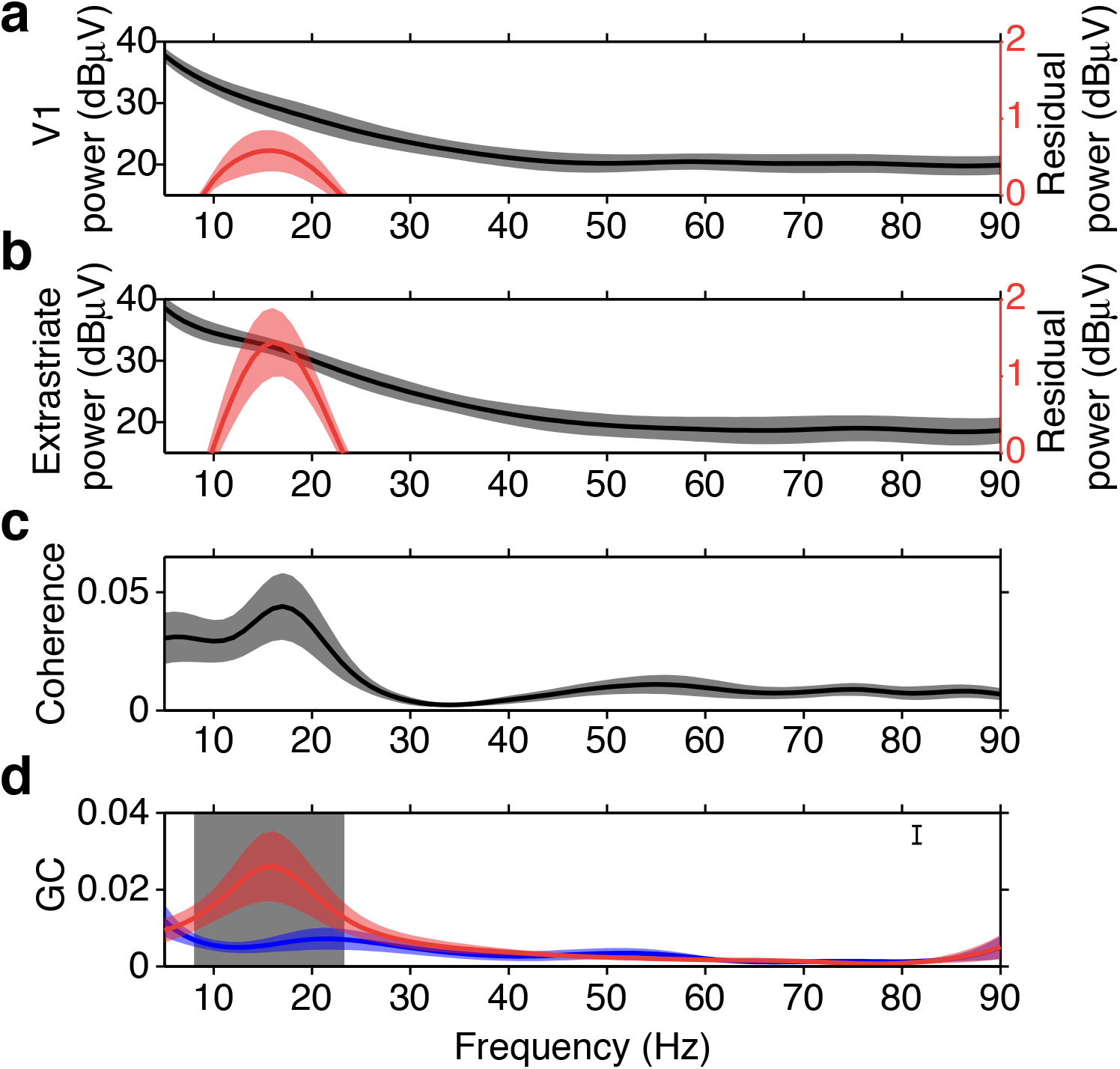
Prestimulus Beta-Frequency Power, Coherence, and sGC Spectra of V1 and V4/TEO LFPs. **a.** Average striate power spectrum over sites (black line ± s.e.m.), and the residual power spectrum after 1/f removal (red line ± s.e.m.) for V1 sites. **b.** Average extrastriate power spectrum over sites and monkeys (black line ± s.e.m.), and the residual power spectrum after 1/f removal (red line ± s.e.m.) for the V4/TEO sites. **c.** Average coherence spectrum over V1-extrastriate site pairs ± s.e.m. for V1-extrastriate pairs. **d.** Average top-down (red line ± s.e.m.), and bottom-up (blue line ± s.e.m.) GC spectra for V1-extrastriate pairs. Shaded grey region denotes the frequencies (8-23 Hz) where top-down and bottom-up sGC were significantly different (p<0.001).

### Prestimulus Oscillatory Synchrony Supports Top-down Signaling to V1

Based on this strong tendency for prestimulus V1 and V4/TEO LFPs to oscillate in the beta frequency range, we hypothesized that beta-frequency synchrony supports top-down signaling from extrastriate cortex to V1. To test this hypothesis, we next computed bottom-up and top-down spectral Granger Causality (sGC) between LFPs in V1 and V4/TEO. The sGC measures statistical causality at spectral frequencies in the Nyquist range, quantifying the degree to which the prediction of a value of one time series can be improved by knowledge of the past values of another time series as a function of frequency.

The mean top-down sGC spectrum showed a peak at 16 Hz, closely matching the frequency of the power peaks and coherence peak, while the mean bottom-up sGC spectrum did not show a beta peak (Figure 2d). A peak in the sGC spectrum indicates narrow-band directed synchrony between the two processes used to derive it. This result supports the hypothesis that beta-frequency synchrony underlies top-down signaling from extrastriate cortex to V1. According to the known relation between coherence and sGC, we inferred that synchronization between V1 and extrastriate areas is dominated by a top-down transfer of information from extrastriate cortex to V1.

To more precisely determine the difference between top-down and bottom-up sGC spectra shown in Fig 2d, we tested for an sGC directional asymmetry at all frequencies between 5 and 90 Hz using a bootstrap resampling approach. We found top-down sGC to be significantly greater than bottom-up sGC only for frequencies between 8 and 23 Hz (grey region in Fig 2d); this range was centered very close to the 16 Hz mean top-down peak (p < 0.001, two-tailed corrected bootstrap resampling test, n = 12). In fact, the probability density of top-down sGC peaks was most prominent in the low-beta-frequency range, with the peak probability density at ~16 Hz being at least 2.5 times larger than at any other frequency examined (Fig 3a). These results point to the presence of top-down physiological signaling from extrastriate low-beta-frequency oscillatory generators to V1 low-beta-frequency generators in subjects awaiting stimulus presentation as they performed the visual pattern discrimination task.

**Figure 3.**
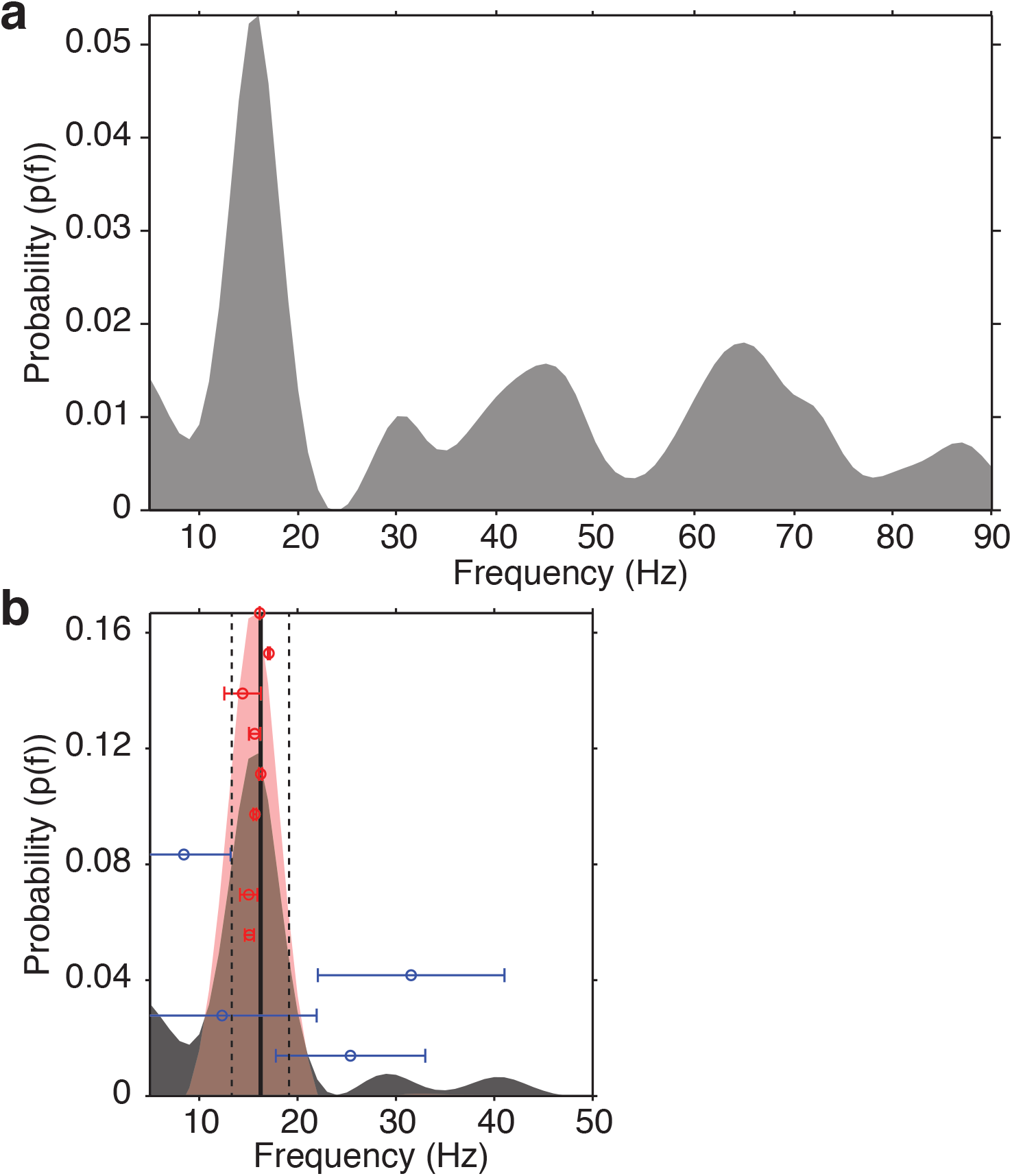
Probability Density of Top-down sGC Peaks. **a.** Probability density as a function of spectral frequency for top-down sGC spectral peaks between 5 and 90 Hz from all V1-extrastriate pairs and bootstrap resamples, showing the greatest probability density at ~16 Hz. **b.** Probability density as a function of frequency for top-down sGC spectral peaks between 5 and 50 Hz from all V1-extrastriate pairs and bootstrap resamples (grey shaded distribution). The mean peak frequency (~16 Hz) across pairs and bootstraps is shown as a solid black vertical line with the 95% confidence interval bounded by vertical dashed lines. The probability density for site pairs having their mean peak frequency inside this 95% confidence interval is shown by the red shaded distribution. The mean peak frequency (± 1 standard deviation) is shown for each of the 12 site pairs, with that for the 8 having their mean peak frequency inside this 95% confidence interval shown by red bars, and that for the other 4 site pairs shown by blue bars.

### Prestimulus Top-down Directed Beta-Frequency Synchrony Predicts Behavioral Context

The evidence for top-down V4/TEO→V1 physiological signaling suggested that the transmission of top-down influences from V4/TEO to V1 may carry task-related behavioral information. We therefore tested the hypothesis that the spatial pattern of top-down beta-frequency sGC contains task-specific behavioral information. To perform this test, we applied a linear Support-Vector-Machine (SVM) classifier, using the top-down beta-frequency peak sGC magnitudes as classification features, in order to predict the task rule (behavioral context) that was in effect. The top-down beta-frequency peak sGC magnitudes used for classification were from the 8 site pairs (out of the 12 possible) that exhibited two important properties: (1) the mean peak frequency was inside the 95% confidence interval of the overall mean of 16 Hz; and (2) the standard deviation of peak frequency was low (Figure 3b, red bars). These 8 site pairs thus showed a consistent top-down directed synchrony in a narrow frequency band around 16 Hz. By contrast, the other 4 site pairs had their mean peak frequency outside the 95% confidence interval and had a large variability of that peak frequency (Figure 3b, blue bars). They thus did not show a consistent top-down directed synchrony in a narrow frequency band. The subsequent analysis focused on these 8 site pairs showing consistent top-down directed synchrony in a narrow frequency band around 16 Hz.

The spatial patterns of these consistent top-down narrow-band beta-frequency sGC peaks, and of their corresponding coherence peaks, are depicted in the maps of Figure 4, where peak coherence is represented by lines connecting V4/TEO and V1 electrode sites, peak sGC is represented by arrows from a V4/TEO site to a V1 site, and line or arrow thickness represents the magnitude of peak coherence or sGC.

**Figure 4.**
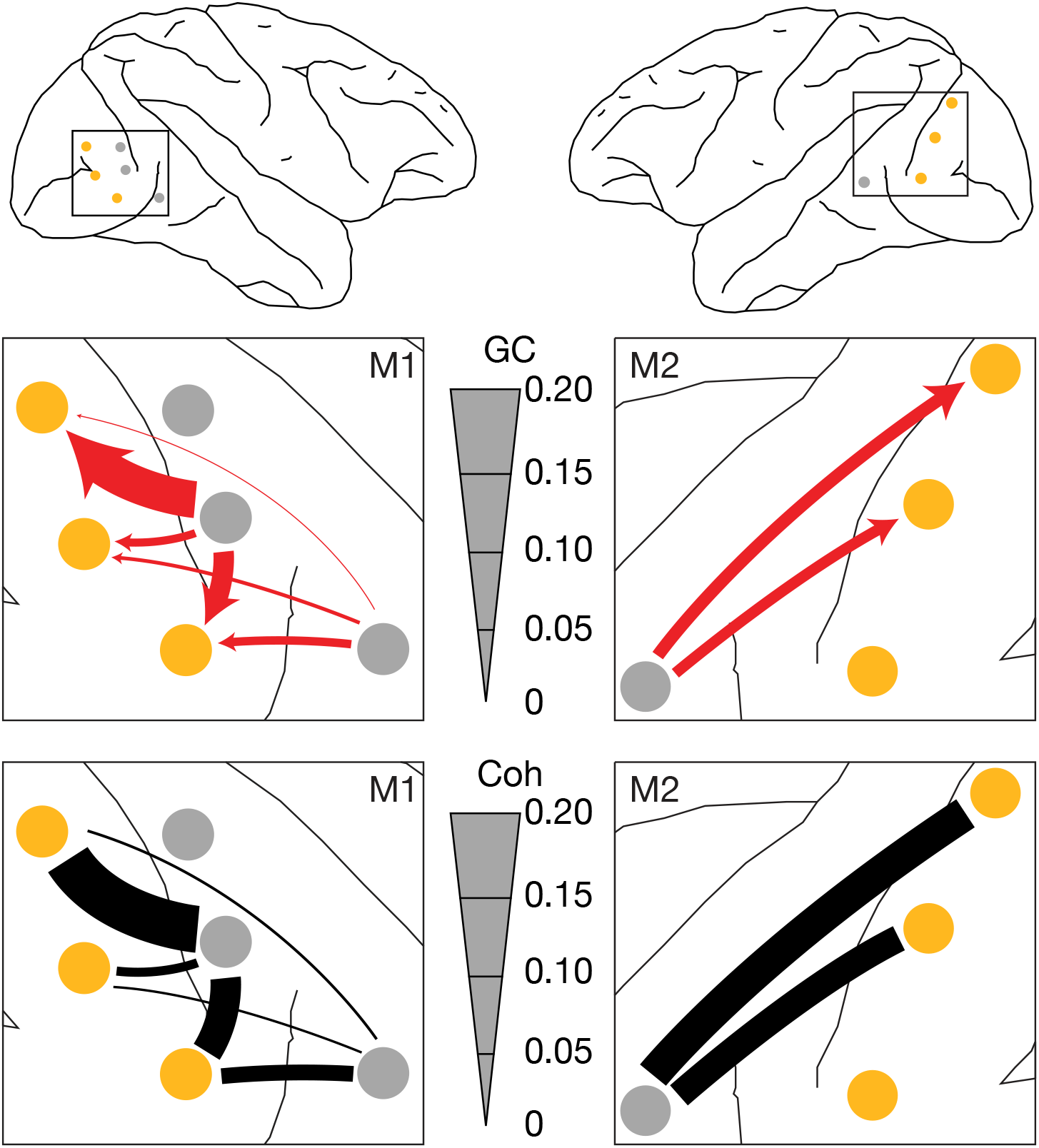
Prestimulus Beta-Frequency Coherence and Top-down sGC Maps. Top: Maps of the recording sites for M1 and M2. V1 electrode locations are marked in gold, and extrastriate (V4 and TEO) locations in grey. Middle: enlarged maps of visual cortex showing top-down sGC at 16 Hz as red arrows for V1-extrastriate pairs having their mean peak frequency inside the 95% confidence interval of Figure 3. Bottom: corresponding maps of coherence for the same site pairs. Thickness of the top-down sGC arrows and coherence bars is proportional to the magnitude of sGC or coherence at 16 Hz.

We found that top-down, but not bottom-up, sGC was highly correlated with coherence (Figure 5). At 16 Hz there was a significant linear correlation (R(6) = 0.90, p < 0.005, Bonferroni corrected) between top-down sGC and coherence values,and top-down sGC explained 81% of the variance in coherence (Figure 5a). By contrast, bottom-up sGC and coherence were not significantly correlated (R(6) = 0.55, p = 0.315, Bonferroni corrected). We also computed the correlation between sGC and coherence at 16 Hz after first aligning the mean sGC and coherence magnitudes and standardizing the variance of each pair over bootstraps (Figure 5b). Even after this normalization, the fraction of the coherence variance explained by the top-down sGC of the 8 site pairs having consistent top-down narrow-beta-band directed synchrony (48%) greatly exceeded that explained by the bottom-up sGC of these site pairs (16%), and even more greatly exceeded that explained by the top-down sGC of the other (inconsistent) 4 site pairs (4%).

**Figure 5.**
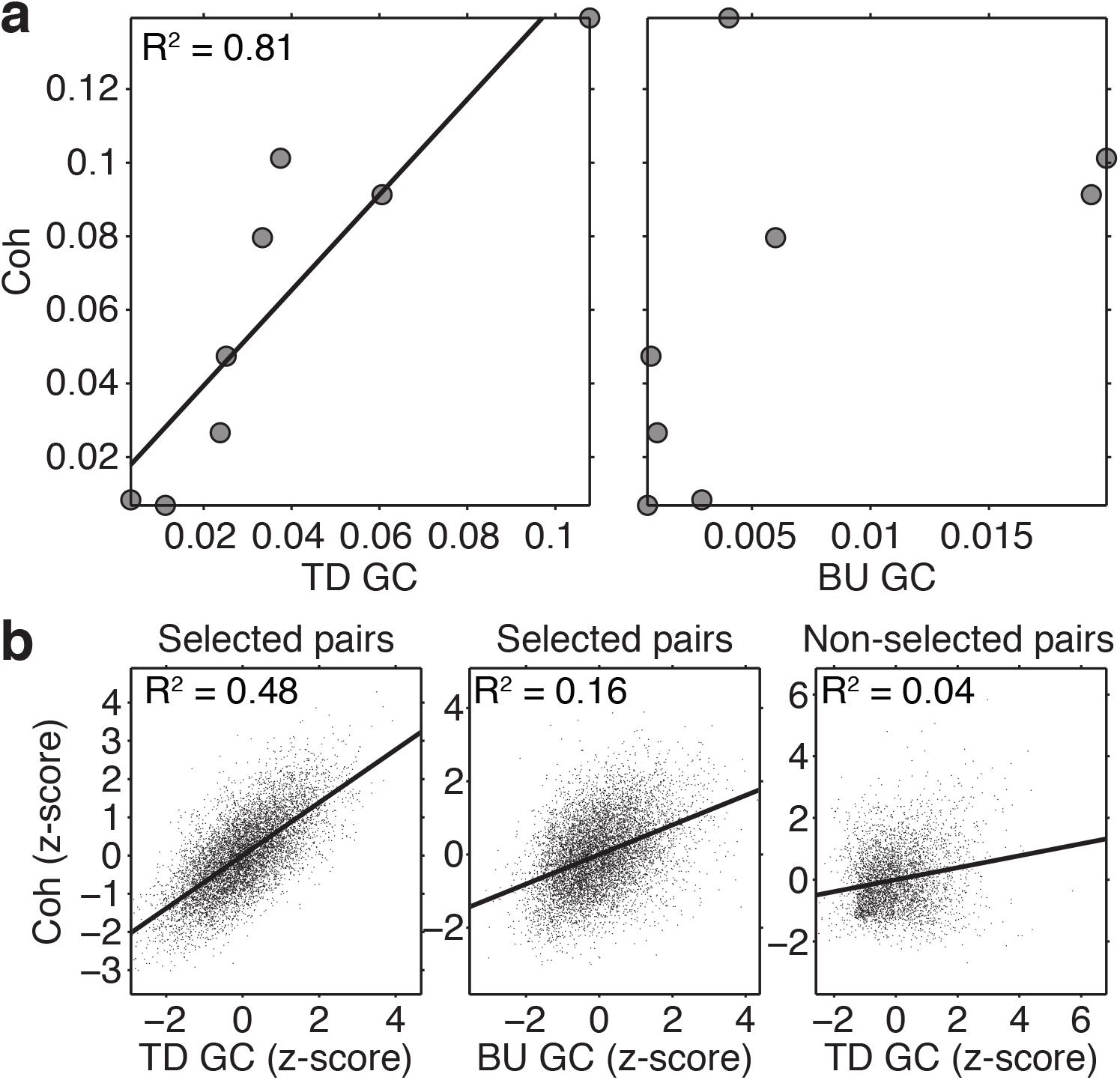
Correlation Between Coherence and sGC. **a.** Correlation between coherence and top-down (left) and bottom-up (right) sGC at 16 Hz across V1-extrastriate site pairs. Coherence was significantly correlated with top-down sGC (R(6) = 0.90, p = 0.005, Bonferroni corrected), but not with bottom-up sGC. **b.** Correlation between normalized coherence and top-down (left), and bottom-up (middle) sGC at 16 Hz averaged over the bootstrap resamples of all 8 V1-extrastriate pairs, having their mean peak frequency inside the 95% confidence interval of Figure 3. The correlation between coherence and top-down and bottom-up sGC explained 48% and 16% of the coherence variance, respectively. Correlation between coherence and top-down (right) sGC at 16 Hz for the other 4 site pairs explained only 4% of the coherence variance.

Support-Vector-Machine (SVM) pattern classification, with the set of top-down narrow-beta-band peak sGC values (taken from the 8 consistent site pairs) as classification features, provided evidence that top-down beta-frequency sGC encodes the task rule (Figure 6). The classification accuracy was 76% for M1, which was significantly greater than the 50% chance classification level (t(10177) = 2.05, p = 0.020), and 82% for M2, which was also significantly greater than the chance level (t(8266) = 2.70, p = 0.004). The results were verified by applying a linear discriminant analysis pattern classifier to the same classification data.

**Figure 6.**
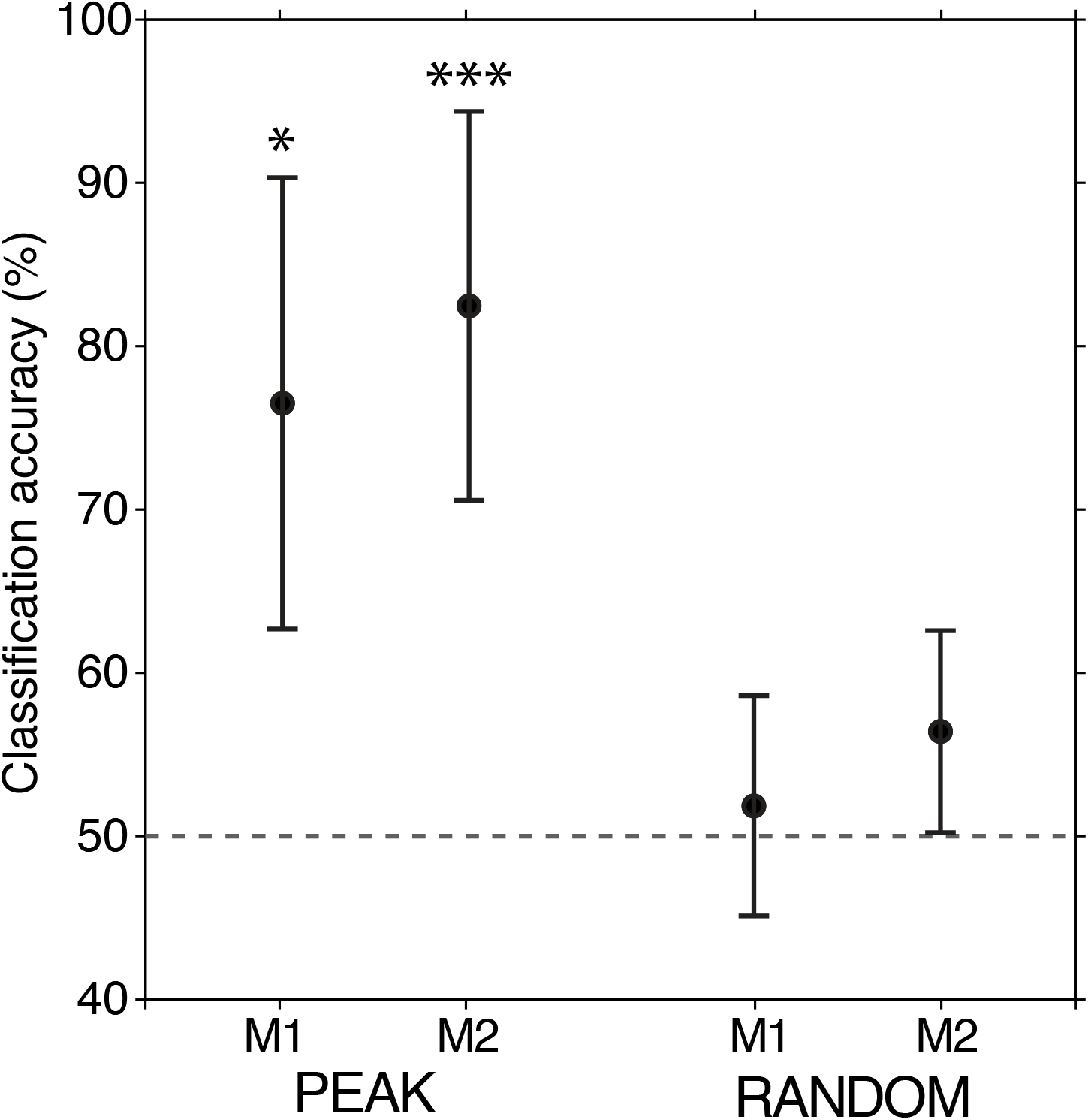
SVM Classification of Task Contingency. Classification accuracy ± s.e.m. based on V1-extrastriate top-down narrow-beta-band sGC peak magnitudes (left group) and randomly selected sGC magnitudes between 5 and 50 Hz (right group). The top-down sGC peak-based classifier significantly exceeded chance (dashed line) for M1 (76%, t(10177) = 2.05, p = 0.020) and M2 (82%, (t(8266) = 2.70, p = 0.004). The SVM classifiers based on randomly selected magnitudes were near the chance level (50%).

To ensure that these results were specifically due to a difference in beta-frequency spectral peak magnitudes between task contingencies, we performed an additional statistical contrast where the classification was based on the top-down sGC magnitude at frequencies randomly selected between 5 and 50 Hz. This resulted in classification accuracies that were near the 50% chance level (Figure 6: M1: 51%, t(10177) = 0.21, p = 0.42); M2: 56%, t(8266) = 1.12, p = 0.132)), thus demonstrating that significant classification performance critically depends on the peak frequencies in a narrow beta-frequency band.

To summarize, these results demonstrate that a top-down narrow-beta-band synchrony directed to V1 from V4/TEO exists in a brief prestimulus window in monkeys highly trained to perform a visuomotor pattern discrimination task, and that this top-down narrow-beta-band directed synchrony predicts the task rule that is in effect. Since task rule determines the behavioral context under which the monkey was performing, the results indicate that behavioral context is conveyed to V1 from extrastriate cortex by top-down beta oscillatory signaling.

## Discussion

We report that prestimulus top-down beta-frequency directed synchrony in visual cortex discriminates the task rule (behavioral context) that governs correct behavioral performance. Our results indicate that task-related behavioral context is conveyed by endogenous top-down neural influences from extrastriate visual cortex (V4/TEO) to primary visual cortex (V1). Extrastriate visual cortex itself likely receives contextual information via influences from higher level areas located in frontal and parietal cortex, via known anatomical projections (Bressler et al. 2008; Markov et al. 2013). Overall, our findings support a cortical model in which contextual information about the behavioral significance of expected stimuli is propagated to V1 by a cascade of top-down influences flowing down a cortical hierarchy (Bressler and Richter 2015).

This main finding is based on the observation that before stimulus presentation in a visuomotor pattern discrimination task top-down directed synchrony is found in a narrow frequency band around 16 Hz for a majority of the site pairs examined. This result suggests the existence of an anticipatory network of visual cortical neuronal populations that are phase-synchronized in a narrow beta-frequency band. The findings of this report are thus consistent with the concept of phase-synchronized large-scale cortical networks that have previously been proposed to operate in cognition (Bressler 1995, 2004, 2008; Bressler and Kelso 2001; Bressler and Tognoli 2006; Bressler et al. 2007; Meehan and Bressler 2012).

The sGC technique (Geweke 1982) that we employed to measure directed synchrony is based on a well-principled and long-established methodology (Bressler and Seth 2013). Although several studies (e.g., von Stein et al. 2000; Saalman et al. 2007) have attempted to infer cortical information transfer from the sign of oscillatory phase difference, it has previously been demonstrated that phase difference may not accurately reflect the direction of influence in cortical circuits (Brovelli et al. 2004; Salazar et al. 2012; Matias et al. 2014).

The linear SVM pattern classification technique was used to discriminate task rule based on top-down beta-frequency directed synchrony entirely from within visual cortex. The classification results were validated by linear discriminant analysis. The classification findings may be considered surprising given that visual cortex is not traditionally considered to process task rules. However, they are consistent with an expanding body of evidence showing that visual processing is contextual (Gilbert and Sigman 2007), and that V1 can be “pre-tuned” in preparation for visual perception (Farber et al. 2015). Only correct trials were considered in this study. Too few incorrect trials were left for analysis after artifact rejection, and so comparison of correct and incorrect performance was not possible.

The top-down extrastriate-to-V1 directed synchrony that we report is in the beta frequency range (~16 Hz), consistent with a growing number of reports relating interareal beta-frequency interactions to endogenous cognitive processing (see reviews in Wang 2010 and Siegel et al. 2012). Although beta oscillations have previously been most associated with somatosensory-motor function (Jenkinson and Brown 2011), accumulating evidence supports the idea that they occur during wait periods when subjects are prepared for a sensory event or for motor behavior (Engel and Fries 2010). We now report that they appear to mediate the expectancy of visual processing as well. Moreover, our finding that top-down beta-frequency synchrony directed to V1 predicts behavioral contingency suggests that beta oscillations may also actively convey endogenous, task-related contextual information to lower-order areas in other (non-visual) sensory systems.

The precise neuronal mechanism by which top-down influences operate in visual cortex is unknown. However, top-down influences from extrastriate cortex may act on V1 inhibitory interneurons to increase their synchrony, and thereby increase their response gain (Lee et al. 2012; Mitchell et al. 2007; Tiesinga et al. 2004). These V1 inhibitory interneurons likely control the infragranular V1 pyramidal neuron, thought to be the principle projection neuron from V1 (Briggs 2014). Modulatory inputs to the interneuron pool are likely to be transmitted by descending fiber pathways, which terminate in both supragranular and infragranular layers. Prominent beta activity has been reported in the infragranular layers of both V1 and extrastriate areas (Buffalo et al. 2011). Thus, in agreement with physiological and modeling studies (Wang 2010; Lee et al. 2012), beta oscillations are a strong candidate for the transmission of top-down influence down the visual hierarchy.

The fact that no stimulus was present during the analysis period of our study suggests the reason why we did not observe prominent gamma-frequency influences. It may be that descending beta and ascending gamma influences can be decoupled in time. If so, top-down beta influences to V1 prior to stimulus onset may modify subsequent V1 stimulus responses. At times, top-down beta influences may also interact directly with stimulus-driven input. Both mechanisms dictate that feedforward stimulus-driven input carried by gamma oscillations is constrained by behavioral context, via descending beta frequency modulation (Bosman et al. 2012; Roberts et al. 2013; Bastos et al. 2015, Richter et al. 2016).

Taken together, our results argue for the idea that extrastriate cortex transmits top-down influences to V1 when well-trained monkeys expect a familiar visual input. We find that these influences (1) depend on synchronous activity between extrastriate and V1 neuron populations, and (2) carry behaviorally relevant task information. The evidence provided here supports the hypothesis that top-down influences from higher areas within the visual cortical hierarchy dynamically constrain lower-level activity in an adaptive, task-specific manner.

## Materials and Methods

### Task

Two well-trained adult rhesus macaque (*Macaca mulatta*) monkeys (M1 and M2) performed a go/no-go visual pattern discrimination task (Figure 1). The stimulus set contained four patterns (each belonging to one of two categories): two “lines” and two “diamonds”. The task rule determined the stimulus-response contingency that governed whether the correct behavioral response to a visual stimulus pattern type (line or diamond) was go or no-go. It was randomly reversed across trial blocks. Control of behavioral context was achieved by (randomly) changing the task rule.

The stimuli were displayed on a screen at a distance of 57 centimeters from the eyes of the subject. Each of the four stimuli consisted of four solid white dots (0.9 degrees visual angle per side), with two of the dots arranged diagonally on opposite corners of an outer square (six degrees visual angle), and the other two dots arranged diagonally on the opposite corners of an inner square (two degrees visual angle) (Figure 1). Line stimuli were patterns where the dots on the outer and inner squares were slanted in the same direction, while diamond stimuli had outer and inner dots slanted in opposing directions. Although the V1 recording sites were expected to have a precise retinotopic relation with the stimulus dots, V1 retinotopic mapping was not available for these monkeys. However, the design of the stimulus ensured that categorization could not be accomplished by observing any single dot, and that the total area, contrast, edge length, and brightness were constant across all stimulus types.

The mean level of correct performance was 92.4 +/− 3.9% for M1 (18 sessions, minimum 84%, maximum 97%) and 95.7 +/− 3.3% for M2 (19 sessions, minimum 89%, maximum 99%). Each trial was initiated when the monkey engaged a lever with the dominant hand. When the lever was depressed and maintained in the depressed position, the trial commenced. After initiation of the trial by the lever press, there was a random period of 200 - 1215 ms before the appearance of the visual stimulus. The visual stimulus was displayed for 100 ms followed by a 400 ms window during which the monkey was required to release the lever on go trials, or maintain lever pressure on no-go trials. Correct go responses were rewarded.

### Electrophysiological Recordings

In both M1 and M2, local field potentials (LFPs), which index the local synaptic activity of the neuronal population at a recording site (Lopes da Silva 2013), were differentially recorded from bipolar Teflon-coated platinum-iridium microelectrodes (more advanced tip near the boundary between the gray and white matter, less advanced tip at the pial surface) chronically implanted at up to 16 cortical sites in the hemisphere contralateral to the dominant hand (for further details see Ledberg et al. 2007). The bipolar microelectrodes were composed of 0.125 mm diameter wires having 2.5 mm tip separation. Electrode positions were verified in one monkey (M1) by both postmortem visual inspection and magnetic resonance imaging. New to the present study, recordings were from areas in the ventral visual stream corresponding to V1, V4, and the temporal occipital area (TEO). Recording sites posterior to the lunate sulcus corresponded to V1, while sites within the prelunate gyrus (V4) and posterior inferotemporal cortex (TEO) were designated as extrastriate cortex. Recordings from M1 were from three V1 recording sites, and three extrastriate recording sites. Of these three extrastriate sites, two were in area V4 and one was in TEO. All recording sites were posterior to the posterior middle temporal sulcus. Recordings from M2 were from three V1 recording sites, and one extrastriate site lying in area V4. The LFP from each bipolar recording electrode was differentially recorded, amplified, and band-passed filtered (-6 dB at 1 and 100 Hz, 6 dB per octave falloff) using a Grass model P511J amplifier, and digitized (12 bits/sample at 200 samples/sec). Differential recording reduced the common contributions to the two electrode tips by more than 10000 times, thus excluding propagated fields from more than a few millimeters away and localizing activity to the tissue between the tips of the bipolar electrode. All experiments were performed by Dr. Richard Nakamura at the Laboratory of Neuropsychology of the National Institute for Mental Health. Animal care was in accordance with institutional guidelines at the time. Surgical methods were as previously described (Ledberg et al. 2007).

The data used in this report were recorded during multiple daily sessions spanning several months, and have not previously been studied. One session was recorded from each monkey per day with a typical recording session composed of 1000 trials. The study employed 18 and 19 sessions from M1 and M2, respectively. Visual and automated artifact rejection reduced the number of correct trials (including both go and no-go responses) available for spectral analysis to 10178 and 8267 for M1 and M2, respectively. An analysis of interactions between other cortical areas in the same monkeys was previously published (Brovelli et al. 2004). However, the current report represents a new analysis of interactions between visual cortical areas. Since interactions between extrastriate cortex and V1 are known to be hierarchical, this study used these regions to investigate cortical top-down and bottom-up influences.

Data acquisition began 85 msec before stimulus onset. Data were not recorded immediately following the lever press due to data storage limitations, and so it was not possible to analyze the temporal evolution of changes taking place between the lever press and the stimulus. Neural activity evoked by the stimulus was absent during the prestimulus period (Ledberg et al. 2007).

### Spectral Analysis

Short-window autoregressive (AR) spectral analysis was performed on all available prestimulus (85 ms) LFP time series data. AR spectral analysis involves application of Fourier-based techniques to an AR model rather than directly to the LFP data. These techniques were utilized instead of direct-data Fourier-based techniques since the spectral resolution of the latter is not sufficient for the short time window analyzed (Nalatore and Rangarajan 2009), whereas the spectral resolution and minimal frequency that may be resolved by the AR approach are not limited by the data period that is analyzed (Cohen 2014; Ozaki 2012). Despite this advantage, AR models might be unstable at low frequencies and near the Nyquist frequency, and so we limited analysis to frequencies between 5 and 90 Hz, based on the 200 Hz sampling frequency of the data. To ensure that each trial of local field potential data could be considered a realization of a zero-mean stochastic process, as required by the AR modeling procedure, the ensemble average was subtracted from each trial for each recording site included in the model (Ding et al. 2006). A model order of 10 was used based on the Akaike Information Criterion (AIC) and previous determination that this value is optimal for this type of data (Brovelli et al. 2004). Extensive testing revealed that the spectral results of this study were not sensitive to the choice of 10 as the model order. In fact, recomputing AR models, with model order varying from 5 to 15, produced spectral peaks having the same peak frequency but differing in peak width (increasing width with decreasing model order).

Subtracting the mean value of the trial ensemble from each trial of the prestimulus V4/TEO and V1 local field potential time series allowed the LFPs to be treated as stochastic processes with stationary mean and variance (Bressler and Seth 2011). We constructed two AR models (restricted and unrestricted) each for each pair of LFPs, represented in the following as variables *X* and *Y*. (Definitions are given for AR models of *X*. Similar definitions may be given for *Y*.) First, the restricted AR model of *X* is defined as:

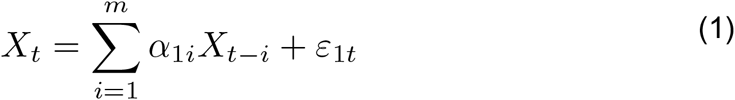

where *X*_*t*_ represents the present value of *X*, *X*_*t*-*i*_ represents past values of *X*, *α*_1*i*_ are the model coefficients, *m* is the model order, and *ε*_1*t*_ is the restricted residual error. Second, *X* may also be represented by the unrestricted AR model:

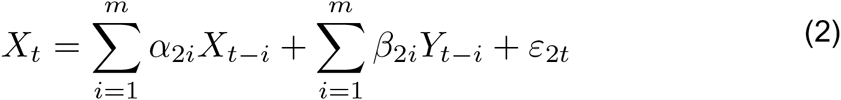

where *Y*_*t-i*_ represents past values of *Y*, *α*_2*i*_ and *β*_2*i*_ are model coefficients, and *ε*_2*t*_ is the unrestricted residual error. Spectral power, coherence, and Granger Causality were computed by well-established methods (Geweke 1982; Ding et al. 2006). For a value of spectral Granger Causality (sGC) to be significant requires that the unrestricted residual error variance be significantly less than the restricted residual error variance. This requirement also controls for the fact that the number of parameters is greater in the unrestricted than the restricted model.

The expression for sGC has a natural interpretation as the fraction of the total power of *X* that is predicted by *Y.* It is expressed as a ratio, where the numerator represents the total power of *X* at a given frequency *ω*, and the denominator represents the intrinsic power, i.e. the power not predicted by *Y.* If the intrinsic power equals the total power, it means that *Y* provides no additional predictability of *X* (in the unrestricted model) above that provided solely by the past of *X* alone (in the restricted model). In the restricted AR model (Equation 1), the causal influence from *Y* to *X* must be zero. The causal power is the amount of additional predictability provided by *Y* in the unrestricted AR model (Equation 2).

sGC can also be viewed as a directional decomposition of neuronal synchronization via its relationship with coherence. The total interdependence is the sum of: 1) the sGC from stochastic process X to stochastic process Y; 2) the sGC from Y to X; and 3) the instantaneous causality, which accounts for instantaneous correlation between X and Y, as would be caused by a mutual simultaneous input to X and Y (see Ding et al., 2006).

To facilitate statistical analysis, site pair identification, and pattern classification, AR spectral estimates were computed on 1000 bootstrap resamples of the data (Efron and Tibshirani 1994), and then averaged over sites (for power spectra) or site pairs (for coherence and sGC spectra). The resulting mean of the resampled spectra is equivalent to the spectrum that would result from an AR model fit over all trials.

To determine frequencies where the top-down and bottom-up spectra significantly differed, a directional asymmetry analysis was performed according to the following procedure (Richter et al. 2016): 1000 bootstrap spectra gave rise to 1000 difference spectra computed (over the entire spectrum from 5 to 90 Hz) as the top-down (V4/TEO to V1) GC spectrum minus the bottom-up (V1 to V4/TEO) GC spectrum for each bootstrap resample. The standard error of the directional asymmetry was then computed from the bootstraps via the following equation (Efron and Tibshirani 1994):

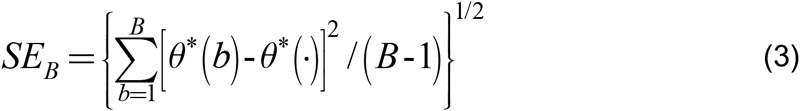

where *B* is the number of bootstraps, in this case 1000, *θ*\*(*b*) is the statistic of interest computed on each bootstrap *b,* and *θ*\*(·) is the mean of all *θ*\*(*b*).

To correct for multiple comparisons across frequencies (Richter et al. 2016), the maximal absolute deviation of each bootstrap difference spectrum from the mean of all bootstrap difference spectra was calculated for each frequency. This resulted in the following modification to Equation 3:

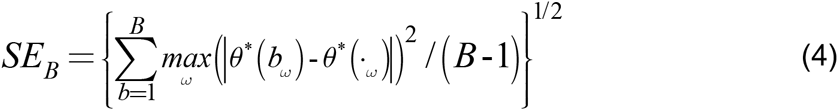

where *ω* indexes each frequency of the spectrum. The resulting standard error was then the maximal standard error that could have been generated across all frequencies and was thus the omnibus standard error (Westfall and Young 1993; Nichols and Holmes 2002; Holmes et al. 1996). Using this standard error, a confidence interval of the mean corresponding to a two-tailed test of p<0.001 was derived as:

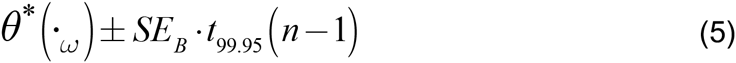

where *n* is the number of trials, and the standard error is multiplied by the t-value with *df* = *n-1*, at the 99.95^th^ percentile of student’s t-distribution. Statistical significance was then assessed at those frequencies where the confidence interval did not contain zero. The frequencies where top-down and bottom-up sGC spectra were significantly different (p<0.001) are indicated by the shaded grey region in Figure 2d, indicating that significant directional asymmetry (top-down sGC > bottom-up sGC) only exists in a limited portion of the spectrum around 16 Hz.

### Removal of 1/f Background Component

The V1 and V4/TEO power spectra had a large 1/f background component that masked the beta oscillatory component (He 2014). To observe the oscillatory component, it was necessary to remove this background component. Power in microvolts was first converted to dB, and then the data were linearized with a logarithmic transform and a line was fit to the data via robust regression between 5 and 30 Hz, using the Welsh weighting function. The resulting regression coefficients were used to specify an exponential function, which was subtracted from the non-log transformed data. The residual is plotted in Figure 2a,b as the de-noised power spectra. The fitting region of 5 – 30 Hz was roughly centered on the dominant top-down beta peak frequency of 16 Hz, such that the robust fit would maximally expose deviations from linearity at this frequency after the log-transform. A wider fitting region would result in greater error due to the contribution of deviations from other possible spectral concentrations, such as in the gamma or delta bands. This also explains why the resulting residual spectrum appears clipped at zero, which is due to the power after the subtraction of the estimated 1/f background component becoming less than zero as a result of error at the left and right extrema of the fit.

### Identification of Top-down Spectral Peaks

A total of twelve V4/TEO-V1 pairs of the recording sites in M1 and M2 were possible. Our goal was to identify the site pairs which had a significant top-down (V4/TEO to V1) spectral peak in the range of frequencies showing significant directional asymmetry (as depicted by the grey region on the mean (across-pair) spectra of Figure 2d). To accomplish this aim, top-down sGC spectra were first computed over the 1000 bootstrap resamples (taken from all trials). Spectral peaks between 5 and 90 Hz were then identified in each bootstrap resample (using the findpeaksG.m Matlab function written by T.C. O’Haver) for all site pairs in the study. Each peak was then tested for significance at the p < 0.05 level against a null distribution.

The null distribution was created as follows: 1) for one null resample, trial order was randomized so that for each AR model the order of trials for site 1 was random with respect to site 2; 2) AR models were fit, and sGC spectra derived; 3) the significance threshold was defined as the maximum value across all frequencies of all top-down sGC spectra; and 4) steps 1-3 were repeated 1000 times to create the null distribution. This procedure controlled for the possibility of spurious results due to multiple comparisons across frequencies and pairs.

The empirical probability density for significant top-down sGC spectral peaks, computed as a function of frequency between 5 and 90 Hz from all site pairs and bootstrap resamples, and fit via a smoothing spline (with a smoothing parameter of 0.075), is presented in Figure 3a. The largest concentration of significant top-down peaks was observed at approximately 16 Hz. To focus analysis on this largest concentration, we computed the empirical probability density by selecting the top-down spectral peak closest to the ~16 Hz peak for all site pairs and bootstrap resamples (shown in Figure 3b). To isolate site pairs with a consistent top-down spectral peak concentration sufficiently near the average (~16 Hz), we computed the mean peak frequency over pairs for each bootstrap (solid vertical line in Figure 3b) and derived its standard error via Equation 3. This allowed us to compute the 95% confidence interval via Equation 5 (vertical dashed lines in Figure 3b). The site pairs having their mean peak frequency inside this 95% confidence interval were considered to have a top-down spectral peak sufficiently close to the peak significant directional asymmetry, indicating that they may be members of a synchronized oscillatory network.

To ensure that the top-down sGC peaks were not driven by differences in signal to noise ratio, we computed the spectral asymmetry for each pair as the difference between top-down sGC and bottom-up sGC, and compared this difference to the same quantity computed on the time reversed data (one time series reversed). As proposed by Haufe et al. (2012), if the difference between the net flow and reverse-net flow is significantly different from zero, then the causal relation is not rejected as spurious, since spurious contributions will not reverse. The test involved computing a single-sample t-test (*df* = 10) for each pair between the net flow difference (net flow – reverse-net flow), and zero, with the two-tailed p-value for all pairs being significant at p<0.05 after Bonferroni correction for multiple comparisons.

### Pattern Classification

A linear Support-Vector-Machine (SVM) was implemented via libSVM (Chang and Lin 2011) to attempt to classify the spatial patterns of prestimulus top-down sGC according to which stimulus-response contingency (task rule) was in effect (Cortes and Vapnik 1995). The go response was the correct response to a line stimulus for contingency 1 trials (no-go response to diamond stimuli), whereas the go response was the correct response to a diamond stimulus for contingency 2 trials (no-go response to line stimuli). The spatial patterns used to train the SVM corresponded to the set of significant magnitudes of the top-down sGC pairs shown by arrows in Figure 4. The machine learning process progressed, individually for M1 and M2, as follows:

1. The trial data for each contingency were randomly split in half into testing and training sets.
2. A bootstrap resample was drawn for the training and testing sets of each contingency, AR models were fit, and top-down sGC spectra were derived. This procedure was repeated 200 times, resulting in 200 exemplars of the top-down sGC pattern, for both training and testing sets of each contingency.
3. The SVM pattern classification feature for each site pair in each of the 200 bootstrap resamples was taken as the magnitude of the closest top-down sGC peak to 16 Hz. If all pairs did not exhibit a peak for a particular bootstrap resample, that resample was deleted. Deletion was a rare event that occurred for less than ½ a percent of all bootstrap resamples (M1: 0.32 %, M2: 0.041). The resulting top-down sGC values were normalized (converted to z-scores) so that each feature (top-down sGC magnitude) had a mean of zero, and unit variance across training and testing sets of both contingencies. Thus the data were not disturbed relative to each condition (training and testing set, and contingency), but the SVM features were balanced (Juszczak et al. 2002).
4. The classification model was created by fitting the linear SVM, with a cost parameter of 1, to the data of the testing set.
5. The classification model from 4) was validated by application to the training data, resulting in a scalar classification accuracy.

These steps together, comprising a delete-d jackknife cross-validation procedure, were repeated 5000 times, producing a mean classification accuracy, and an estimate of its standard error.

Statistical evaluation of the SVM results was performed via application of delete-d jackknife cross-validation (Efron and Tibshirani 1994). Delete-d jackknife cross-validation entails training (model fitting) on a portion of the total data (*n − d*), and then testing the model accuracy on the remaining *d* samples. This is an attractive approach for the current problem since sGC is biased as a function of the number of trials. Thus, by selecting *d* to be half the data, the bootstrap sGC will have equal trial numbers yielding equivalent bias across training and testing groups and between contingencies. Since the number of contingency 1 and contingency 2 trials was not exactly equal, each bootstrap resample used selections with an n equal to the contingency with the lowest number of trials. The delete-d jackknife is computed according to the following formula:

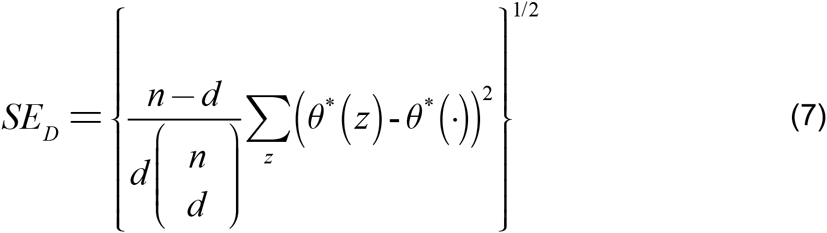

where *n* is the number of trials, 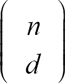 is the total number of possible selections of *d* elements that can be made from *n, θ*\*(*z*) is the statistic computed on each of the possible subsets, and *θ*\*(·) is the mean of that statistic over subsets. In the current case of approximately 10 000 trials per monkey, the number of possible subsets is effectively infinite. However, Shao (1989) has demonstrated that a Monte Carlo approach known as the jackknife-sampling variance estimator (JSVE) well approximates the variance of the estimator. The standard error based on the JSVE method is estimated as:

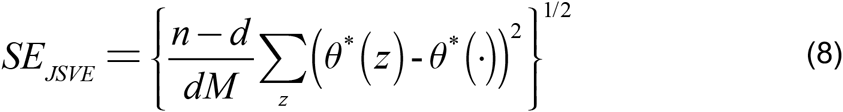

where *M* is a random number of subsets of *d* selected from the total possible. Shao (1989) shows that even when *M* is dramatically smaller than the total number of possible selections, the JSVE still outperforms a number of estimators, such as the bootstrap. Supplementary Fig 1 shows that in the current data the standard error estimate quickly reaches an asymptote as *M* exceeds 250 subsamples, and remains very stable as it approaches the 5 000 subsamples used to estimate the standard error. In addition, the estimation error falls to negligible levels as M increases beyond. Using the standard error derived via the JSVE procedure, a single-sample t-test was conducted to determine if the mean classification accuracy differed from the chance level of 50%.

## Acknowledgments

The authors gratefully acknowledge Richard Nakamura for his role in designing the experimental paradigm and performing the electrophysiological recordings. This work was supported by the U.S. National Institute of Mental Health under grant MH062404.

---

**Supplementary Figure 1.**
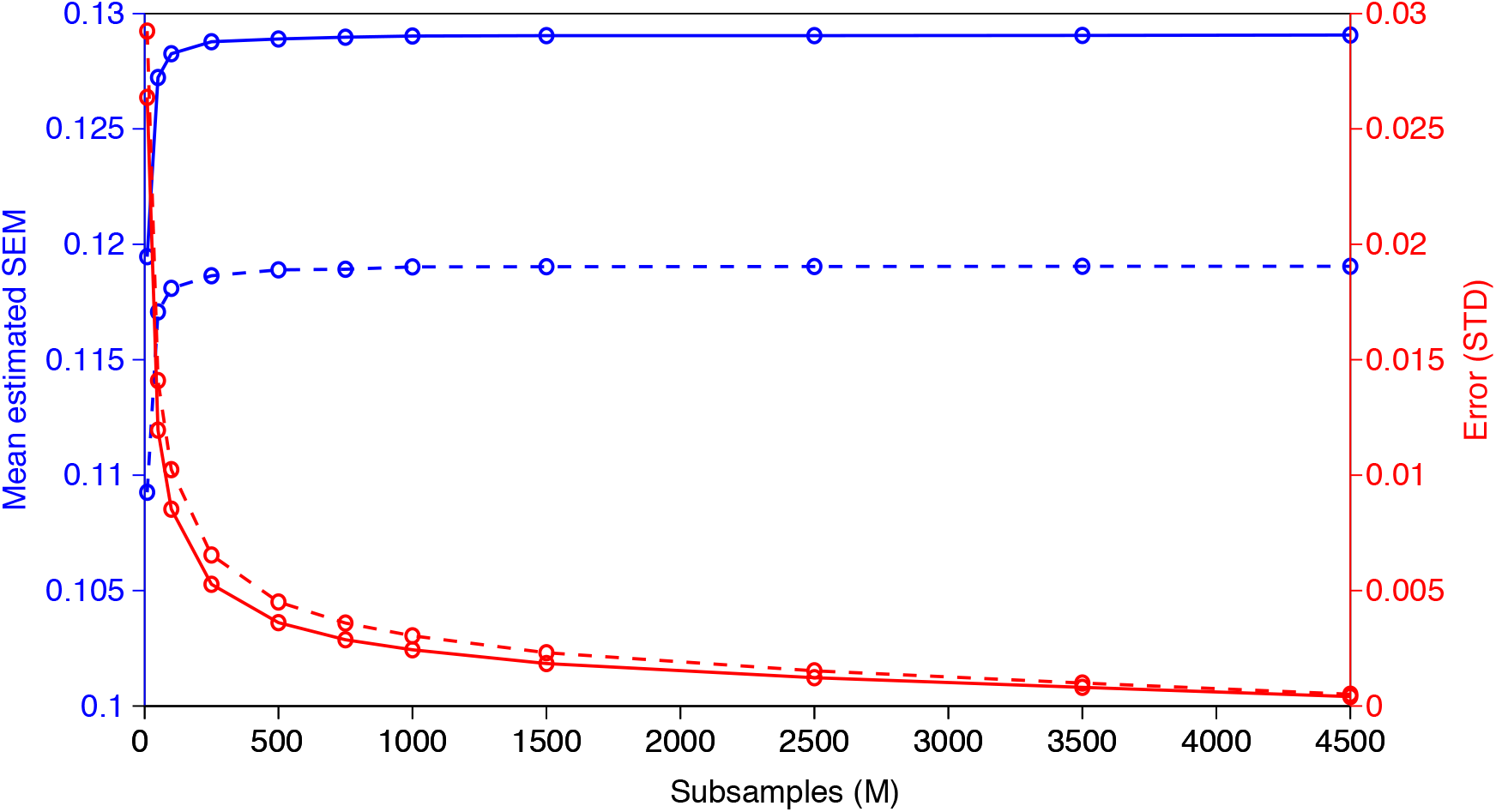
Average Estimated Standard Error of the Mean and Estimator Variance as a Function of Subsamples. Average estimated standard error for M1 (solid blue curve) and M2 (dashed blue curve) as a function of subsamples. The estimate stabilizes above a subsample size of 250. The variance of the estimator for M1 (solid red curve) and M2 (dashed red curve), estimated over 10000 random subsamples for each level of M, monotonically decreases with subsample size quickly become negligible as M increases.

**Supplementary Figure 2.**
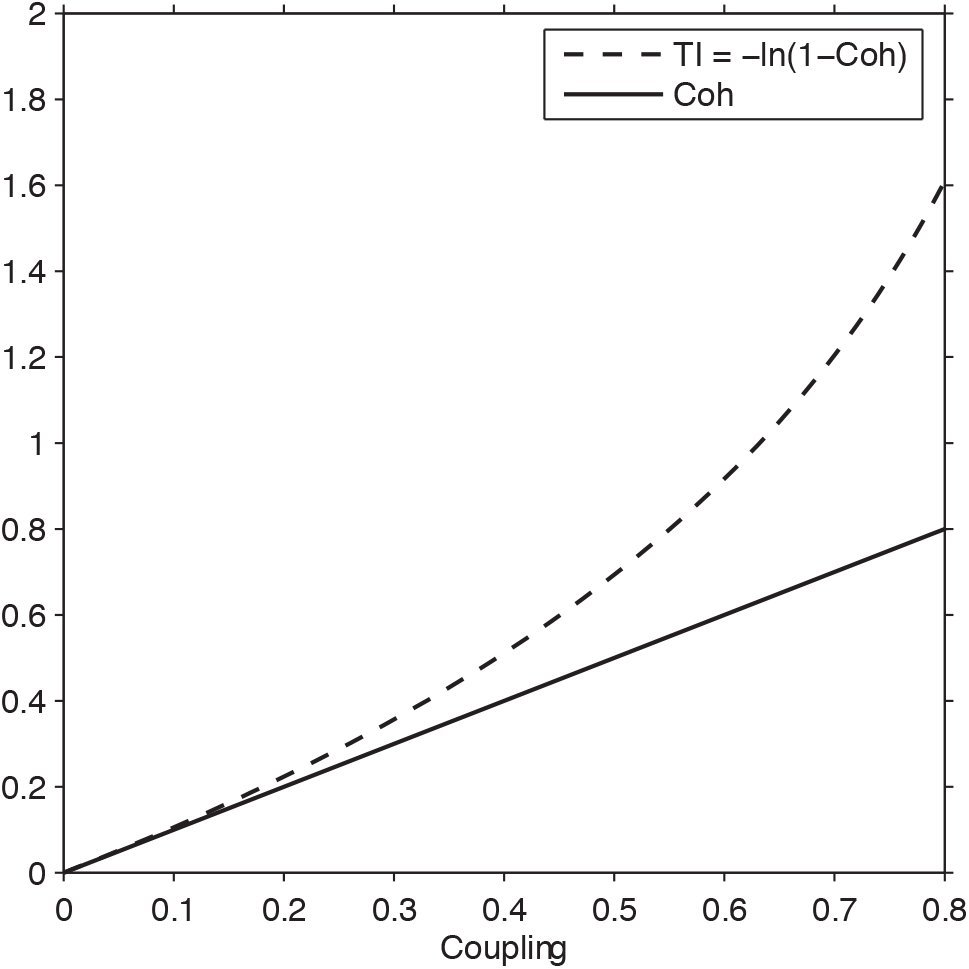
Coherence and Total Interdependence as a Function of Coupling Strength. Coherence and Total Interdependence are strongly correlated for physiologically realistic levels of coherence (~< 0.2). Values are computed based on the coherence, which was derived using unit power for both simulated signals and a coupling term (numerator of the coherence equation) varied between 0 and 0.8.

